# Host infection dynamics and disease induced mortality modify species contributions to the environmental reservoir

**DOI:** 10.1101/2022.09.20.508714

**Authors:** Nichole A. Laggan, Katy L. Parise, J. Paul White, Heather M. Kaarakka, Jennifer A. Redell, John E. DePue, William H. Scullon, Joseph Kath, Jeffrey T. Foster, A. Marm Kilpatrick, Kate E. Langwig, Joseph R. Hoyt

## Abstract

Environmental pathogen reservoirs exist for many globally important diseases and can fuel epidemics, influence pathogen evolution, and increase the threat of host extinction. Species composition can be an important factor that shapes reservoir dynamics and ultimately determines the outcome of a disease outbreak. However, disease induced mortality can change species communities, indicating that species responsible for environmental reservoir maintenance may change over time. Here we examine reservoir dynamics of *Pseudogymnoascus destructans,* the fungal pathogen that causes white-nose syndrome in bats. We quantified changes in pathogen shedding, infection prevalence and intensity, host abundance, and the subsequent propagule pressure imposed by each species over time. We find that highly shedding species are important during pathogen invasion, but contribute less over time to environmental contamination as they also suffer the greatest declines. Less infected species remain more abundant, resulting in equivalent or higher propagule pressure. More broadly, we demonstrate that high infection intensity and subsequent mortality during disease progression can reduce the contributions of high shedding species to long-term pathogen maintenance.

## Introduction

Emerging infectious diseases threaten efforts to conserve global biodiversity (Daszak et al. 2000, Taylor et al. 2001, Jones et al. 2008, Fisher et al. 2012). In some disease systems, pathogens may survive for long periods of time in the environment in the absence of a living host (Turner et al. 2016, Plummer et al. 2018, Islam et al. 2020). Pathogen persistence in the environment allows for transmission independent of infected hosts, can exacerbate disease impacts, and increase the risk of host extinction (de Castro and Bolker 2005, Mitchell et al. 2008, Almberg et al. 2011, Hoyt et al. 2020). However, pathogen contamination in the environment is not homogenous; rather, variation in the amount of pathogen in the environmental reservoir is likely driven by a complex process of pathogen shedding from hosts within the community leading to subsequent transmission events.

Infected hosts can vary in the amount of pathogen they shed into the environment with some hosts producing disproportionately high amounts of pathogen, independent of direct host contacts (Sheth et al. 2006, Chase-Topping et al. 2008, Lawley et al. 2008, Direnzo et al. 2014). Research has shown that variation in host shedding can be driven by differences in behavior (Godfrey 2013, Rushmore et al. 2013, VanderWaal and Ezenwa 2016), innate susceptibility (Searle et al. 2011, Gervasi et al. 2013), space use (Brooks-Pollock et al. 2014), and infection severity (Lloyd-Smith et al. 2005, Munywoki et al. 2015). In multi-host disease systems, variation in pathogen shedding, produced through community composition and species abundance, can play a key role in transmission dynamics (Kilpatrick et al. 2006, Esteban et al. 2009, Paull et al. 2012).

Host abundance within a community can interact with host shedding to moderate transmission from a particular species (Lloyd-Smith et al. 2005, Paull et al. 2012, Kilonzo et al. 2013). Species that have low rates of shedding, but are highly abundant, may contribute more to transmission than might be expected based on the per capita amount of pathogen they shed (Peterson and McKenzie 2014, Scheele et al. 2017). Conversely, a species that has high rates of shedding or is highly infectious, but at low abundance, may contribute less to transmission than other species (Lloyd-Smith et al. 2005, Kilpatrick et al. 2006). In addition, for some wildlife infectious diseases, infectiousness, shedding, and impacts are positively correlated (Langwig et al. 2016, Brannelly et al. 2020), such that hosts initially important for disease transmission, suffer from high disease-related mortality and become less important contributors to pathogen maintenance over time (Brannelly et al. 2018). Variation in pathogen shedding and how it influences disease dynamics is important for many disease systems (Sheth et al. 2006, Chase-Topping et al. 2008, Henaux and Samuel 2011, Brooks-Pollock et al. 2014, Direnzo et al. 2014, Slater et al. 2016), but how differences in host shedding scale to a community-level and influence the environmental reservoir are rarely linked together.

White-nose syndrome (WNS) is an emerging infectious disease caused by the fungal pathogen *Pseudogymnoascus destructans* (Lorch et al. 2011, Warnecke et al. 2012), that has had devastating effects on bat populations (Langwig et al. 2012, Frick et al. 2015, Langwig et al. 2016). White-nose syndrome exhibits seasonal infection dynamics that are driven by the environmental reservoir and host-pathogen ecology (Langwig et al. 2015a, Hoyt et al. 2021, Langwig et al. 2021, Kailing et al. 2023). *Pseudogymnoascus destructans* can persist for long periods of time in the environment, which results in widespread infection when hosts return to hibernacula (subterranean sites where bats hibernate in the winter) in the fall (Lorch et al. 2013, Hoyt et al. 2015, Langwig et al. 2015a, Campbell et al. 2019, Hoyt et al. 2020, Hicks et al. 2021). During this time, susceptible bats become infected or reinfected by *P. destructans* when they come into contact with the environmental reservoir upon entering hibernacula (Langwig et al. 2015a). Over the winter hibernation period, *P. destructans* grows into the skin tissue, causing deleterious physiological changes, including increased arousals from hibernation, weight loss, dehydration, and often death (cumulative 95-99% declines) (Warnecke et al. 2013, Verant et al. 2014, McGuire et al. 2017, Hoyt et al. 2021). During hibernation, susceptible bat species vary greatly in their infection intensities and three species have suffered declines that exceed 95% (Langwig et al. 2012, Langwig et al. 2016, Hoyt et al. 2020, Hoyt et al. 2021). Species abundance also varies greatly within bat communities and during the epizootic (Langwig et al. 2012, Frick et al. 2015). Together, differences in pathogen shedding and species abundance may influence pathogen contamination in the environment.

The amount of *P. destructans* in the environment has been shown to increase after the first year of invasion (Hoyt et al. 2020) and contamination of the environmental reservoir has been linked to increased pathogen prevalence and loads for bats (Hoyt et al. 2020, Hoyt et al. 2021). As a result, bat mortality also increases with higher levels of environmental contamination (Hoyt et al. 2018, Hoyt et al. 2020, Hicks et al. 2021, Hoyt et al. 2023). However, the establishment of the environmental pathogen reservoir in these multi-host communities remains an important knowledge gap.

Environmental transmission is an important driver of infectious disease dynamics and understanding the factors that lead to pathogen establishment in the environment is crucial for disease control and prevention. Here we use a unique dataset that encompasses the stages of *P. destructans* invasion and establishment to capture pathogen accumulation in the environmental reservoir across 19 sites in the Midwestern United States. Using these data, we explored potential differences in pathogen shedding among species. We also assessed the relationship between bat infection intensity and the amount of pathogen shed into the environment by each species present in the community. We hypothesized that bat species abundance would also play a key role in site-level contamination and environmental reservoir establishment, so we also explored how differential pathogen shedding among species and their abundance influences the propagule pressure within communities.

## Methods

### Sample collection and quantification

We quantified *P. destructans* fungal loads on bats and from hibernacula substrate throughout bat hibernation sites in the Midwestern, United States. Samples were collected from 19 sites in Wisconsin, Illinois, and Michigan and each site included three years of pathogen data from invasion to establishment, which was collected over a seven-year period (Appendix S1: Table S1). Hibernacula were visited twice yearly, once during early hibernation (November to December) and once during late hibernation (March to April) to capture differences in infection dynamics and environmental contamination at the beginning and end of hibernation. During each visit, we counted the total number of bats within each site by species (*Eptesicus fuscus* (Big brown bat)*, Myotis lucifugus* (Little brown bat)*, Myotis septentrionalis* (Northern long-eared bat) and *Perimyotis subflavus* (Tricolored bat)) (Appendix S1: Figure S1). We collected epidermal swab samples from bats within sites to quantify bat infection intensity (quantities of fungal DNA) and determine infection prevalence (Appendix S1: Table S2). Samples were collected using previously established protocols that consisted of rubbing a polyester swab dipped in sterile water over the muzzle and forearm of the bat five times (Langwig et al. 2015a, Hoyt et al. 2016).

To measure the amount of *P. destructans* shed into the environment we also collected environmental substrate swabs from beneath or directly adjacent to each hibernating bat (on hibernacula walls and ceilings) where they are in direct contact with the substrate. Samples collected in close proximity to bats have shown to be strongly tied to the infection intensity of the bat (Langwig et al. 2015b, Hoyt et al. 2020). To capture independent site-level *P. destructans* environmental contamination, we collected swab samples as described above, but collected from the environment in locations more than two meters away from roosting bats, in areas where bats might roost. These samples were used to estimate the reservoir contamination across the site without targeting substrate used by specific bat species. These environmental samples were taken by swabbing an area of substrate equal to the length of a bat’s forearm (36-40 mm) five times back and forth, as described previously (Langwig et al. 2015b). We preserved *P. destructans* DNA samples by storing all swabs in salt preservation buffer (RNAlater; Thermo Fisher Scientific) directly after collection. DNA was extracted from all samples with a modified Qiagen DNeasy Blood & Tissue Kit (Frick et al. 2015, Langwig et al. 2015b). The presence and quantity of *P. destructans* was determined by quantitative Polymerase Chain Reaction (qPCR) (Muller et al. 2013).

To verify that fungal loads measured using qPCR accurately reflected viable fungal spores in the environment, that are able to infect a host, we collected additional substrate swab samples from a subset of locations that were paired with substrate swabs used for qPCR. These samples were cultured by streaking the substrate swab across a plate containing Sabouraud Dextrose Agar treated with chloramphenicol and gentamicin. The plates were stored at 4 °C and colony forming units (CFU’s) of *P. destructans* were quantified within six weeks of initial inoculation. We paired substrate samples analyzed using qPCR to determine log10 *P. destructans* loads for comparison with colony forming units obtained from culture samples to validate viability. There was a significant relationship between quantity of *P. destructans* DNA measured through qPCR and the number of CFU’s (Appendix S1: Figure S2). This suggests that qPCR was a valid method to estimate the amount of *P. destructans* in the environment and supports that qPCR results are reflective of the number of infectious propagules in the environment and not relic DNA (Appendix S1: Figure S2).

All research was approved through Institutional Animal Care and Use Committee protocols: Virginia Polytechnic Institute: 17-180; University of California, Santa Cruz: Kilpm1705; Wisconsin Endangered/Threatened Species Permit 882 & 886; Michigan Department of Natural Resources permit SC-1651; Illinois Endangered/ Threatened Species Permit 5015, 2582 and Scientific Collections permit NH20.5888; US Fish and Wildlife Service Threatened & Endangered Species Permit TE64081B-1.

### Data analysis

We separated invasion stage into two distinct categories: “invasion” which included the first year the pathogen arrived, as described previously (Langwig et al. 2015b, Hoyt et al. 2020), and “establishment” which included the second and third years of *P. destructans* presence in a site when bat species declines begin to occur, which corresponds to the epidemic stage as has been previously noted (Langwig et al. 2015b, Hoyt et al. 2020). We used these stages to capture pathogen shedding into the environment before and after pathogen accumulation in the environment occurred (invasion and establishment, respectively) and to examine the dynamic changes between stages of pathogen invasion and establishment.

We first examined the presence and quantity of *P. destructans* on each bat species and the amount each bat species shed into the environment. We used mixed effects models with log_10_ environmental fungal load collected under each bat as our response variable, bat species as our predictor, and site as a random effect for both invasion stages. Tables are reported for model output by including *M. septentrionalis* as the reference level for the invasion stage and *M. lucifugus* during the established stage. These were chosen because they had the highest levels of shedding in the respective stages and demonstrate contrasts to all other species. We similarly compared infection intensity among species using the same model as described above, but with log_10_ fungal loads on bats as the response variable. For this analysis we reported the table for the model output with *E. fuscus* as the reference level because it has the lowest levels of infection intensity and demonstrates contrasts to the other species. We examined differences in infection prevalence by bat species using a generalized linear mixed effects model with a binomial distribution and a logit link with species as our predictor, bat infection status (0|1) as our response, and site as a random effect. To examine within host variation in pathogen shedding, we calculated the coefficient of variation of pathogen shedding for each species in both invasion stages.

To examine how differences in bat infection intensity contributed to contamination of the environment under each individual, we used a linear mixed effects model to explore the relationship between infection intensity of each bat and the amount of pathogen shed into the environment under each individual. In this analysis, we used paired log_10_ environmental *P. destructans* loads under a bat as our response variable with log_10_ bat fungal loads interacting with species as our predictor and site as a random effect. We combined the invasion and established stages since the amount of pathogen shed into the environment was hypothesized to be a product of how infected the host was, and therefore, comparable across years. We report the output of our model by using estimated marginal means of linear trends to highlight the support for the relationship between bat fungal loads and environmental fungal loads beneath bats for all species and to display multiple species slope contrasts. To examine within host variation in infection intensity, we calculated the coefficient of variation for infection intensity for each species in both invasion stages.

We investigated the role of bat species abundance on environmental contamination of *P. destructans*. We first examined the differences among species abundance within sites for each invasion stage by using a linear mixed effects model with species as our predictor and log_10_ population count during early hibernation (before over-winter declines occur) as our response and included site as a random effect. To explore how species abundance influenced the degree of pathogen contamination within sites, we used a linear mixed effects model with log_10_ environmental fungal loads collected greater than two meters from any bat during late hibernation as our response variable and log_10_ average population abundance between established years interacting with species identity as our predictors with site as a random effect.

Finally, we calculated differences in propagule pressure among bat species by multiplying pathogen prevalence by bat species abundance within each site to get the number of infected individuals. We then multiplied the number of infected individuals by the average amount of fungal spores shed into the environment by each species in each site. We analyzed the propagule pressure calculations using a generalized linear mixed effects model with a negative binomial distribution to account for dispersed population abundance counts with an interaction between species as our predictor and propagule pressure as our response for each invasion stage. We performed this analysis for invasion (year 0) and establishment (years 1-2) during late hibernation to investigate how pathogen pressure may differ across stages of pathogen invasion when bats are heavily shedding into the environment. For the invasion year, we report the table for model output with *M. septentrionalis* as the reference level as it has the highest level of propagule pressure compared to the other species. For the established stage we report using estimated marginal means of our generalized linear mixed effects model to highlight contrasts between multiple species. To assess if propagule pressure was a suitable metric for pathogen invasion of the environmental reservoir, we examined the relationship between site-level environmental contamination, as described above, and sum propagule pressure for each site using a linear mixed effects model with site included as a random effect.

All analyses were conducted in R v.4.2.1 (R Core Team 2022). Mixed-effects models were run using the package “lme4” (Bates et al. 2015) except for analyses using a negative binomial distribution which were ran using “glmmTMB” (Brooks et al. 2017). The reported estimated marginal means and estimated marginal means of linear trends were generated using the package “emmeans” (Searle et al. 1980, Lenth 2022).

## Results

We found that pathogen shedding (the amount of pathogen detected under bats) and infection varied both within and among the four species present in the community (Figure 1, Appendix S1: Figure S3). During initial pathogen invasion into bat communities, we found that on average, individual *M. septentrionalis* (Figure 1A; Appendix S1: Table S4; intercept = -3.72 ± 0.33) contributed more pathogen into the environment than *E. fuscus* (coeff = -0.88 ± 0.33, P = 0.009), *M. lucifugus* (coeff= -0.52 ± 0.22, P = 0.02), and *P. subflavus* (coeff =-0.86 ± 0.35, P = 0.02), and also had the greatest within species variability (Appendix S1: Figure S3A). In the established stage (Figure 1B; Appendix S1: Table S5), we found support for shifts in species shedding patterns with higher shedding in *M. lucifugus* (intercept = -3.41 ± 0.12) than *E. fuscus* (coeff = -0.75 ± 0.14, P < 0.0001) and *P. subflavus* (coeff = -0.65 ± 0.13, P < 0.0001), but similar shedding to *M. septentrionalis* (coeff = -0.33 ± 0.24, P = 0.17).

**Figure 1:**
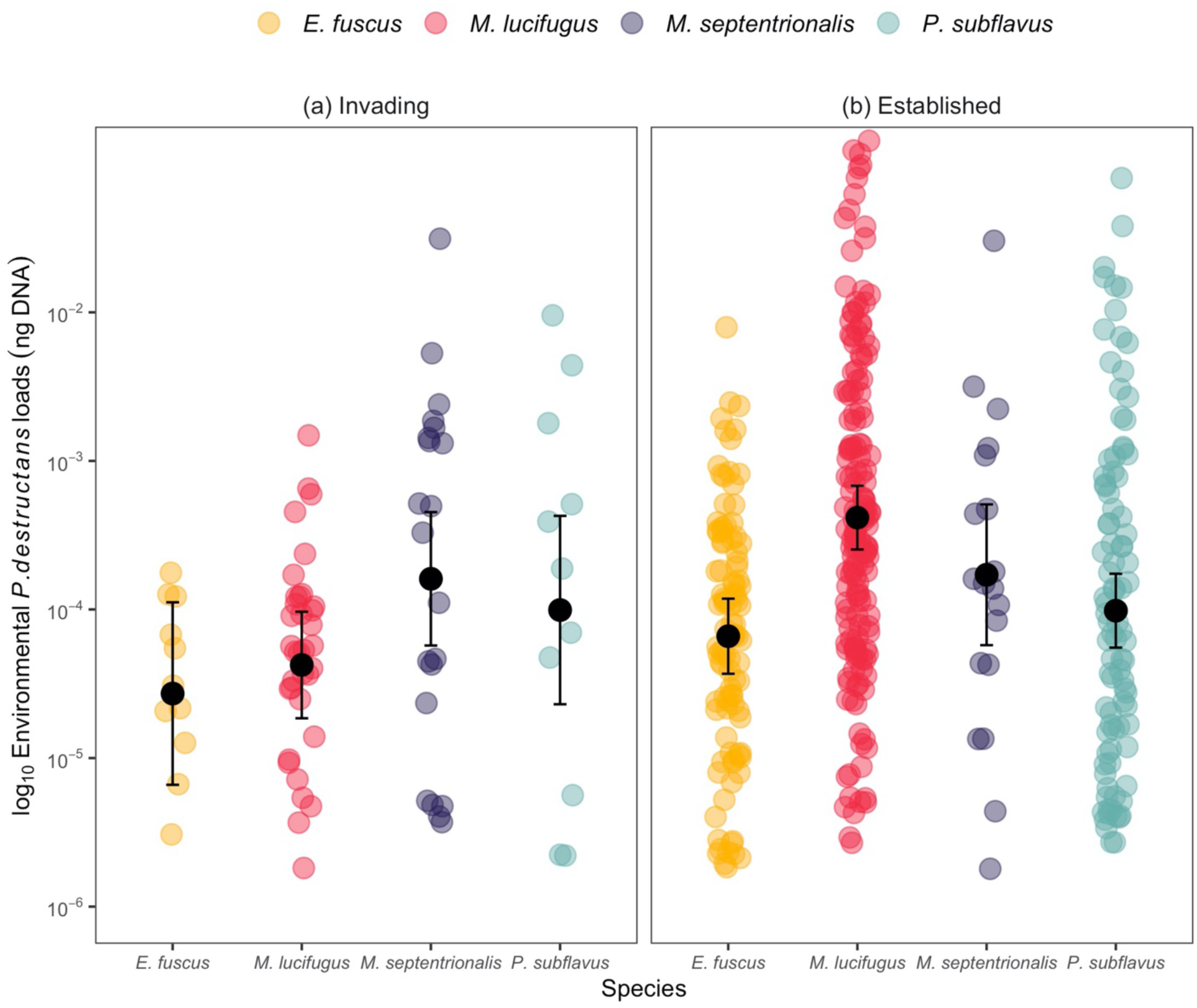
Differences in pathogen shedding into the environment among invading and established invasion stages. log_10_ *P. destructans* environmental loads (ng DNA) among species during pathogen invasion (a) and establishment (b) in late hibernation. Each point represents the log_10_ environmental *P. destructans* loads from under an individual bat. Black points represent the estimated mean and bars indicate ± standard error for each species. Samples collected that were beyond the limits of detection were set to 10^-5.75^ log_10_ *P. destructans* loads (ng DNA).

Our results showed consistent support that host infection intensity predicted the amount of pathogen shed into the environment for all species (Figure 2C; Appendix S1: Table S8; host infection and shedding relationship: *M. lucifugus* slope = 0.38 ± 0.05, P < 0.0001; *E. fuscus* slope = 0.33 ± 0.07, P < 0.0001; *M. septentrionalis* slope = 0.41 ± 0.10, P < 0.0001; *P. subflavus* slope = 0.33 ± 0.08, P = 0.0001). Importantly, we found no support for differences among species in the slope between environmental pathogen shedding and host infection intensity (Figure 2C; Appendix S1: Table S8; all P > 0.05), suggesting that differences in shedding were primarily driven by the infection intensity of individual bats. *Eptesicus fuscus* (intercept = -3.37 ± 0.16) had significantly lower infection intensity than all other species within the community (Figure 2B; Appendix S1: Table S7; *M. lucifugus* coeff = 1.38 ± 0.15, P < 0.0001; *M. septentrionalis* coeff = 1.08 ± 0.21, P < 0.0001; *P. subflavus* coeff = 1.30 ± 0.17, P < 0.0001), but had the greatest amount of variation for within species infection intensity (Appendix S1: Figure S3B). In addition, we found that *M. lucifugus* (intercept = 2.19 ± 0.27) had the highest infection prevalence, with 87% of hosts infected on average across the invasion and established stages (Figure 2A; Appendix S1: Table S6; *E. fuscus* coeff = -0.79 ± 0.31, P < 0.01; *M. septentrionalis* coeff = -0.83 ± 0.35, P = 0.02; *P. subflavus* coeff = -1.10 ± 0.24, P < 0.0001).

**Figure 2:**
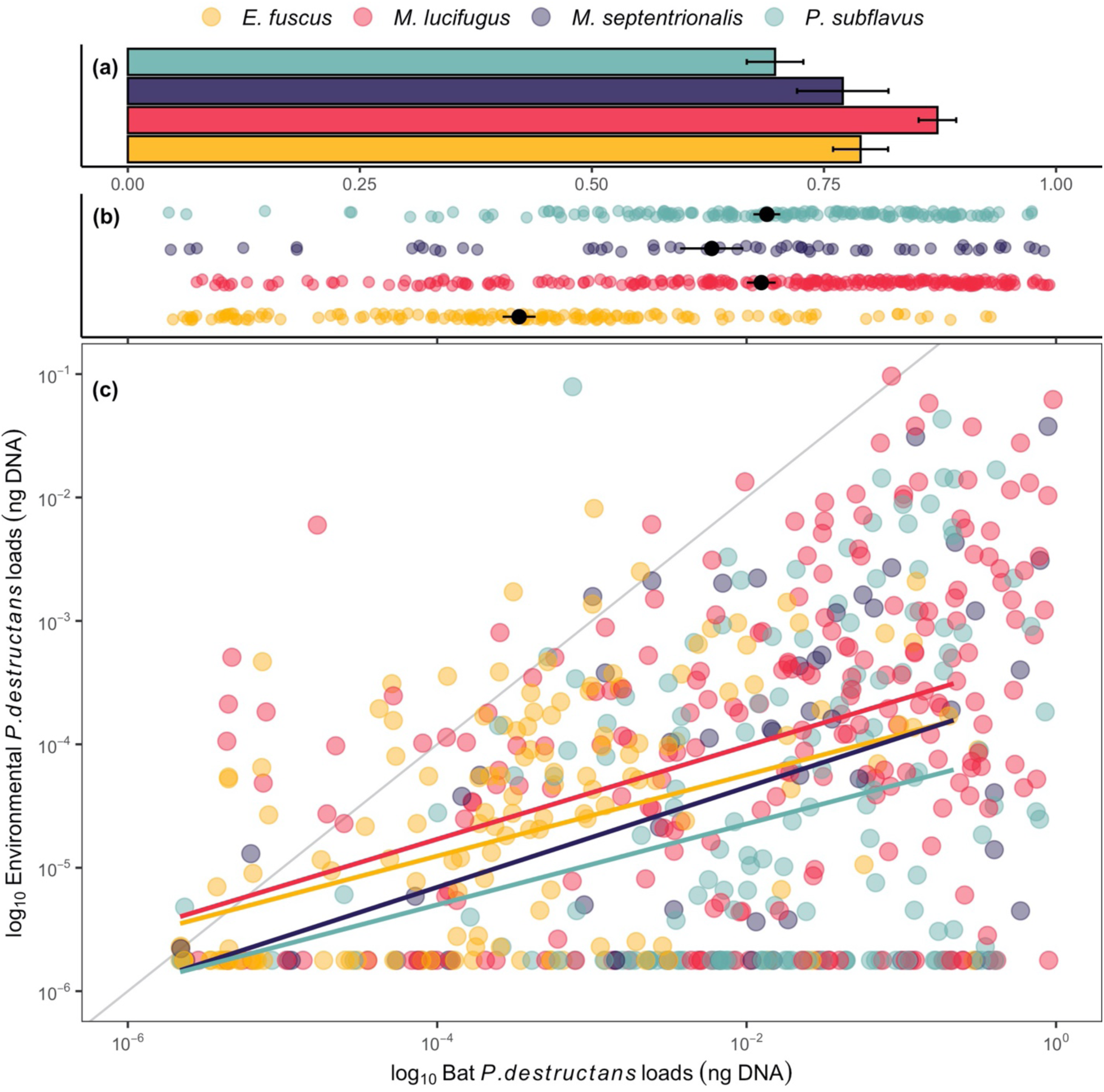
Bat infection prevalence, intensity, and the relationship between host infection and pathogen shedding. For all panels, bat species are displayed by color. Black points indicate mean bat fungal loads by species and bars represent ± standard error. (a) Pathogen prevalence for each species during pathogen invasion in late hibernation. (b) Bat log_10_ *P. destructans* loads (ng DNA) for each species during pathogen invasion in late hibernation. (c) The relationship between log_10_ *P. destructans* loads (ng DNA) on an individual bat and the amount of log_10_ *P. destructans* (ng DNA) in the environment directly underneath individuals during pathogen invasion and establishment in late hibernation. Colored lines show the relationship between log_10_ bat *P. destructans* loads and log_10_ environmental *P. destructans* loads fit with a linear mixed effects model. The gray solid line shows the 1:1 line, and points along this line would indicate that the amount of *P. destructans* on bats was equivalent to the amount shed into the environment. Each individual bat is represented by a point. Samples collected that were beyond the limits of detection were set to 10^-5.75^ log_10_ *P. destructans* loads (ng DNA).

In addition to differences in infection intensity and prevalence, species also varied in abundance, with *M. lucifugus* (intercept = 1.63 ± 0.19) being the most abundant species across sites, with an average population size of 39.81 ± 7.76 before declines from WNS (invasion) (Figure 3; Appendix S1: Table S9; *E. fuscus* coeff = -0.65 ± 0.24, P < 0.01; *M. septentrionalis* coeff = -0.60 ± 0.21, P < 0.01; *P. subflavus* coeff = -0.41 ± 0.23, P < 0.08). During the established stage, while population declines were occurring, *M. lucifugus* (intercept = 1.28 ± 0.13) remained the most abundant species within the community with an average population size of 18.20 ± 7.41 (Figure 3; Appendix S1: Table S10; *E. fuscus* coeff = -0.54 ± 0.16, P < 0.001; *M. septentrionalis* coeff = -0.76 ± 0.14, P < 0.0001; *P. subflavus* coeff = -0.38 ± 0.15, P < 0.01). *M. septentrionalis* had the lowest abundance in the community with an average colony size of 11.22 ± 5.50 individuals before disease impacts and declined to an average of 3.52 ± 4.12 individuals in a site during the established stage (Figure 3; Appendix S1: Table S9; Appendix S1: Table S10). Furthermore, host abundance was important in influencing the degree of site-level environmental pathogen contamination (Appendix S1: Figure S4; Appendix S1: Table S13; *M. lucifugus* slope = 0.25 ± 0.11, P = 0.03).

**Figure 3.**
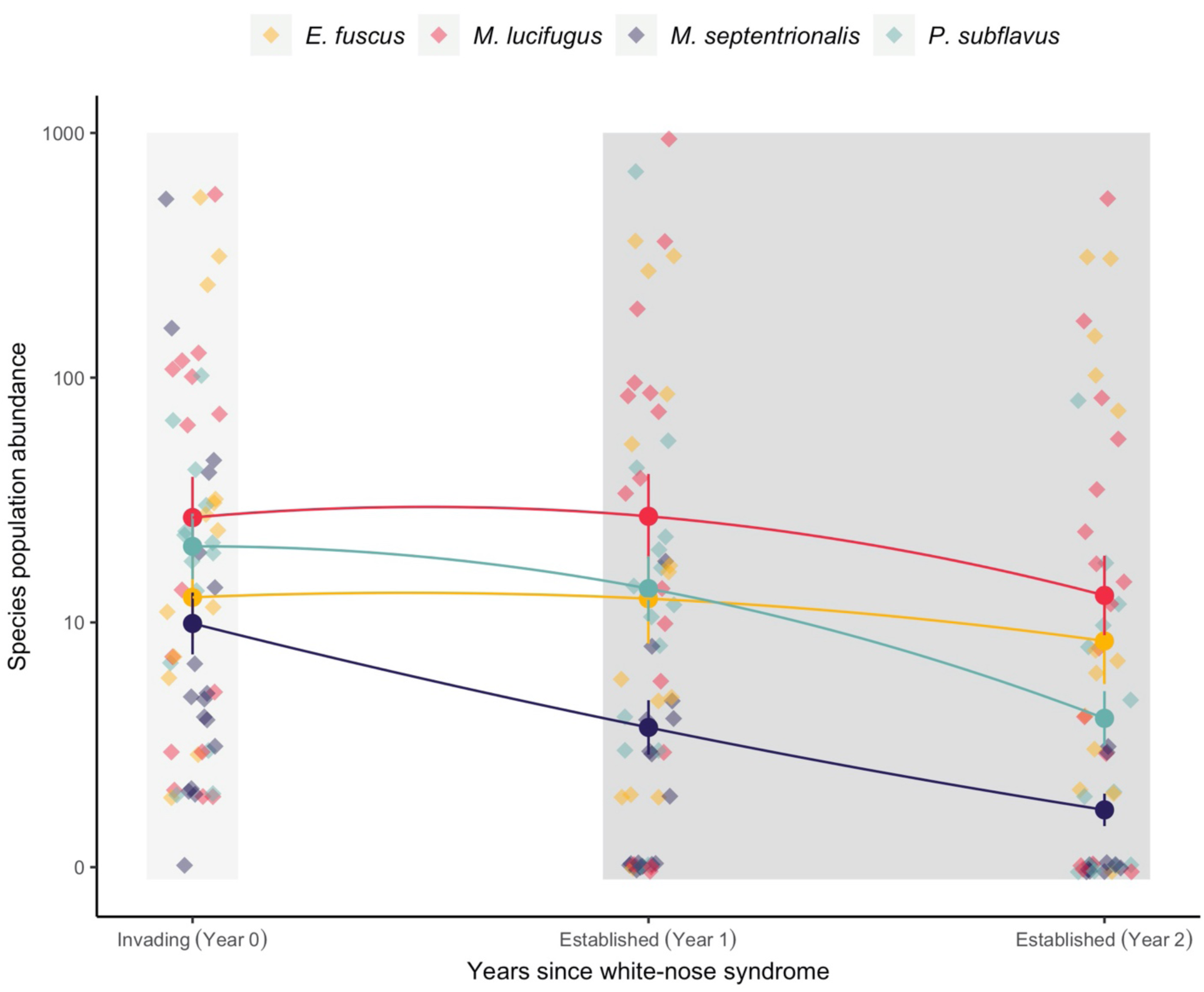
Changes in host abundance between invading and established years. Species within the community are differentiated by color and points represent a species population at an individual site. Species population abundance in the invasion (light gray box) and established years (dark gray box) in late hibernation within each site. Circular points indicate the mean abundance for each year and bars denote ± standard error. For best data visualization, three data points have been excluded from the figure (1,110 *M. lucifugus* and 1,985 *P. subflavus* during the invading stage, and 3,154 *M. lucifugus* during the established stage).

Finally, when we combined infection prevalence, species abundance, and pathogen shedding into the metric of propagule pressure, we found that during the first year of invasion, *M. septentrionalis* (Figure 4A; Appendix S1: Table S11; intercept = 7.44 ± 1.05) had higher propagule pressure than *P. subflavus* (coeff = -3.17 ± 1.25, P = 0.01) and *E. fuscus* (-4.28 ± 1.21, P < 0.001), but not *M. lucifugus* (coeff = 0.87 ± 1.03, P = 0.39). In years following invasion, once *P. destructans* was established, *M. lucifugus* (Figure 4B; Appendix S1: Table S12; coeff from estimated marginal mean = 10.75 ± 0.51) had consistently higher propagule pressure and contributed more pathogen to the environmental reservoir than all other species (Figure 4B; Appendix S1: Table S12; *M. lucifugus*-*M. septentrionalis* contrast = 4.24 ± 0.79, P < 0.001; *M. lucifugus-E. fuscus* contrast = 3.84 ± 0.56, P < 0.0001; M. lucifugus-*P. subflavus* contrast = 2.46 ± 0.54, P = 0.0001). *Myotis septentrionalis* (Figure 4B; Appendix S1: Table S12; coeff from estimated marginal mean = 6.52 ± 0.81) no longer contributed more than *P. subflavus* (*M. septentrionalis*-*P. subflavus* contrast = -1.78 ± 0.85, P = 0.16) or *E. fuscus* (*M. septentrionalis-E. fuscus* contrast = -0.39 ± 0.83, P = 0.97) due to their rarity following disease-induced declines (Figure 4). Mean environmental contamination in areas greater than two meters from bats increased with total propagule pressure calculated at a site level(summed propagule pressure among species), suggesting the measure of propagule pressure accurately captures environmental contamination levels (Figure 4C; relationship between environmental contamination and propagule pressure intercept = -5.48 ± 0.14, slope = 0.12, P = 0.002).

**Figure 4.**
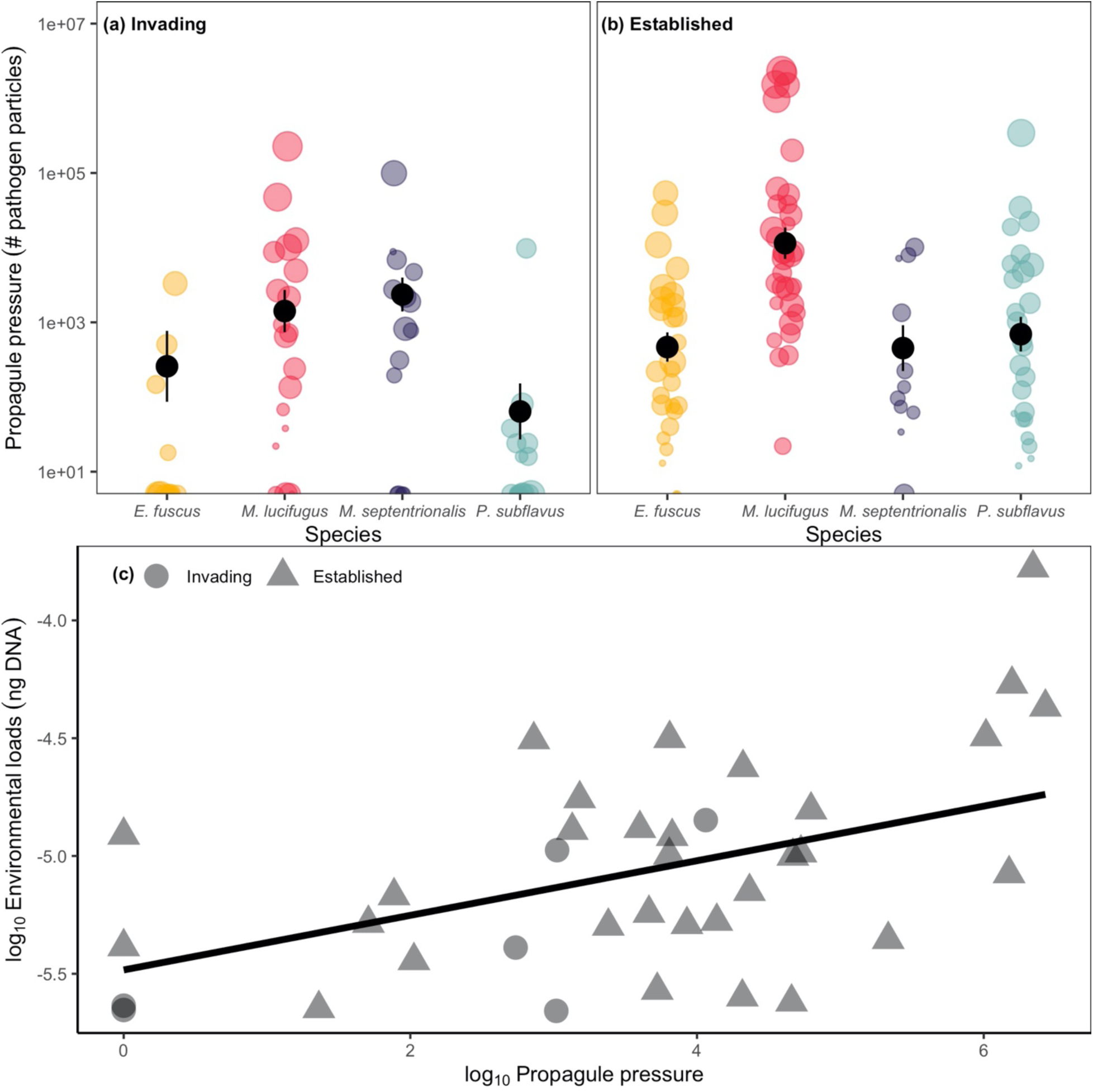
Differences in propagule pressure among species during invading and established stages and the relationship between propagule pressure and environmental contamination. (a-b) Propagule pressure (# of pathogen particles) by species during late hibernation in (a) invading and (b) established stages. Colored points represent populations of species within a site, black points represent the mean and bars indicate ± standard error for each species. Point size is weighted by species population abundance. (c) Relationship between log_10_ propagule pressure and log_10_ mean site-level contamination. Site-level contamination was assessed in samples > 2m away from roosting bats within sites during late hibernation in the established stage. Points indicate the sum propagule pressure at a site for each year and shape denotes invasion stage. The line shows the relationship between log_10_ propagule pressure and log_10_ environmental *P. destructans* loads fit with a linear mixed effects model. The total environmental contamination increased as propagule pressure increased within a site (Environmental contamination: intercept = -5.48 ± 0.14, slope = 0.12, relationship between environmental contamination and propagule pressure P = 0.002).

## Discussion

Variation in pathogen shedding, and subsequent environmental transmission can have dramatic impacts on community-level disease outcomes. Our findings demonstrate how increases in host infection intensity can lead to increased pathogen shedding of *P. destructans* (Figure 2C), which is consistent with research in other systems (Lloyd-Smith et al. 2005, Matthews et al. 2006, Chase-Topping et al. 2008, Direnzo et al. 2014, Munywoki et al. 2015, Maguire et al. 2016, VanderWaal and Ezenwa 2016). In addition, we demonstrate that changes in host abundance and differences in infection prevalence modify species contributions to the environmental reservoir, which varied over time (Figure 2A; Figure 3; Figure 4). Most importantly, while high infection intensity initially leads to high pathogen shedding for one species (Figure 2C), this also leads to elevated mortality (Langwig et al. 2016) and local extirpations (Appendix S1: Figure S1). In contrast, species with lower infection intensity and greater variation within species for infection intensity, such as *E. fuscus*, had reduced impacts (Figure 2B; Appendix S1: Figure S1; Appendix S1: Figure S3B) (Langwig et al. 2016) and may be more important in pathogen maintenance than high shedding species that suffer severe mortality. Therefore, we identified a shift in the species responsible for the greatest environmental reservoir contamination between invasion stages, suggesting that contributions to the reservoir is a dynamic process that varies over time (Figure 4).

The relationship between individual host shedding into the environment and host infection intensity did not differ among species, suggesting that individuals of different species with the same infection intensity are equally efficient at depositing pathogen into the environment (Figure 2C; Appendix S1: Figure S5). Instead, the prevalence and intensity of infection, which did vary among species, influenced how much pathogen was shed on average per individual (Figure 2A; Figure 2B; Figure 2C). Some species such as *E. fuscus* had low-intensity infections despite moderate prevalence, while other species, *M. septentrionalis* and *M. lucifugus*, were much more heavily infected and shed more pathogen into the environment (Figure 2B; Figure 2C). In addition, we saw shifts in host infection intensity between invasion and establishment stages (Figure 2B) leading to shifts in shedding (Figure 1A; Figure 1B). An increase in host infection prevalence between invasion stages has been previously described by Hoyt et al. 2020, where increases in pathogen contamination during the initial invasion stage, results in rapid reinfection and higher exposure doses in subsequent years. This influenced fungal burdens on bats and eventually population level declines (Langwig et al. 2015b, Hoyt et al. 2020, Langwig et al. 2021).

Disease-caused declines in several bat species (Figure 3), which has been shown to be strongly linked with intensity of infection (Langwig et al. 2017, Hopkins et al. 2021), resulted in large changes in species abundance and community composition (Appendix S1: Figure S1). We found support that species abundance predicted site-level contamination (Appendix S1: Figure S4), and this reinforced the need to account for abundance when determining species-level contributions to the environmental reservoir. To combine these factors, we used propagule pressure or “introduction effort”, which is a fundamental ecological metric that determines whether a biological invasion will be successful in reaching establishment (Lockwood et al. 2005), and is especially applicable to emerging infectious diseases. We found that environmental contamination in the sampled locations that were not associated with an individual bat increased as our calculated propagule pressure metric increased (Figure 4C), suggesting that propagule pressure is an accurate way of measuring species contributions to the environmental reservoir.

During the initial invasion of *P. destructans*, both *M. lucifugus* and *M. septentrionalis* had higher propagule pressure than other species present (Figure 4A), which was not apparent through examination of only pathogen shedding (Figure 1A). The high infection intensity of *M. septentrionalis* resulted in a large contribution to the initial establishment of the environmental reservoir despite their low relative abundance (Figure 2B; Figure 3). However, in subsequent years, as this species declined precipitously (Figure 3; Appendix S1: Figure S1), their contribution was greatly reduced and equivalent to less infected species (e.g. *E. fuscus*, Figure 4B).

Our results suggest that host abundance, infection prevalence, and infection intensity (which influences shedding into the environment) are collectively required to describe species contributions to environmental pathogen contamination. For example, while *E. fuscus*, *M. septentrionalis*, and *P. subflavus* eventually contributed similar propagule pressures during the established invasion stage (Figure 4B), this was likely driven by different factors that need to be considered when evaluating their influence on environmental contamination. *Eptesicus fuscus* had low infection intensity and subsequently low declines, but had moderate pathogen prevalence (Figure 2A; Figure 2B) and moderate abundance (Figure 3), which elevated its contribution. *Perimyotis subflavus* had high infection intensity, but reduced prevalence compared to the other heavily infected species (Figure 2A; Figure 2B) and was only present in moderate abundances (Figure 3). Finally, *M. septentrionalis* was heavily infected (Figure 2B), but at low abundance across communities (Figure 3), which reduced its overall importance in years following initial pathogen invasion (Figure 4B). *Myotis lucifugus* maintained high propagule pressure through the invasion and establishment of *P. destructans* (Figure 4A; Figure 4B), suggesting that the presence of *M. lucifugus* within a community will result in rapid establishment of *P. destructans* in the environment, and maintenance of environmental contamination within a site over time (Appendix S1: Figure S4). We also found that within-species variation in infection and pathogen shedding changed across invasion stages by species (Appendix S1: Figure S3) which may be influenced by numerous factors (e.g. timing of infection, microclimate use, etc.) and warrants further investigation. Our results highlight numerous factors that must be considered to evaluate the influence of pathogen shedding on environmental reservoir maintenance, subsequent environmental transmission, and how this can change dynamically across invasion stages.

Understanding environmental reservoir dynamics is crucial for many globally important disease systems. Discerning how variation across species contributes to maintenance of the environmental reservoir is important in determining the extent of epidemics, predicting long-term impacts on host communities, and developing control strategies. We demonstrate that the establishment and maintenance of the environmental reservoir is strongly influenced by variation in pathogen prevalence, infection intensity, and species abundance. Evaluating these effects together through the metric of propagule pressure allowed us to capture which species within the community contributed to pathogen invasion success and ultimately the maintenance of indirect transmission, which is an important driver of infection and mortality (Hoyt et al. 2018). Broadly, our results demonstrate that multiple factors of species variability can scale to influence environmental reservoir dynamics within communities.

## Acknowledgements

The research was funded by the joint NSF-NIH-NIFA Ecology and Evolution of Infectious Disease award DEB-1911853, NSF DEB-1115895, USFWS (F17AP00591), and by the NSF GRFP. We would like to acknowledge E. McMaster, and the numerous other landowners for access to sites, M.J. Kailing for assistance with culture collection and B. Heeringa and D. Kirk for field assistance.

## Author Contributions

NAL and JRH wrote the original draft of the manuscript. JRH, KEL, and AMK designed methodology; NAL, JRH, KEL, AMK, JPW, HMK, JAR, JED, WHS, and JK collected the data; NAL conducted the laboratory experiment; JTF and KLP supervised and performed sample testing; NAL analyzed the data with assistance from KEL and JRH; All authors contributed critically to draft revision.

## References

1. Almberg, E. S., P. C. Cross, C. J. Johnson, D. M. Heisey, and B. J. Richards. 2011. Modeling routes of chronic wasting disease transmission: environmental prion persistence promotes deer population decline and extinction. PloS one 6:e19896.

2. Bates, D., M. Mächler, B. M. Bolker, and S. C. Walker. 2015. Fitting linear mixed-effects models using lme4. Journal of Statistical Software 67:1–48.

3. Brannelly, L. A., H. I. McCallum, L. F. Grogan, C. J. Briggs, M. P. Ribas, M. Hollanders, T. Sasso, M. Familiar López, D. A. Newell, and A. M. Kilpatrick. 2020. Mechanisms underlying host persistence following amphibian disease emergence determine appropriate management strategies. Ecology Letters.

4. Brannelly, L. A., R. J. Webb, D. A. Hunter, N. Clemann, K. Howard, L. F. Skerratt, L. Berger, and B. C. Scheele. 2018. Non-declining amphibians can be important reservoir hosts for amphibian chytrid fungus. Animal Conservation 21:87–180.

5. Brooks, M. E., K. Kristensen, K. J. van Benthem, A. Magnusson, C. W. Berg, A. Nielson, H. J. Skaug, M. Maechler, and B. M. Bolker. 2017. glmmTMB Balances Speed and Flexibility Among Packages for Zero-inflated Generalized Linear Mixed Modeling. The R Journal 9:378–400.

6. Brooks-Pollock, E., G. O. Roberts, and M. J. Keeling. 2014. A dynamic model of bovine tuberculosis spread and control in Great Britain. Nature 511:228–231.

7. Campbell, L. J., D. P. Walsh, D. S. Blehert, and J. M. Lorch. 2019. Long-term survival of Pseudogymnoascus destructans at elevated temperatures. Journal of Wildlife Diseases.

8. Chase-Topping, M., D. Gally, C. Low, L. Matthews, and M. Woolhouse. 2008. Super-shedding and the link between human infection and livestock carriage of Escherichia coli O157. Nat Rev Microbiol 6:904–912.

9. Daszak, P., A. A. Cunningham, and A. D. Hyatt. 2000. Emerging infectious diseases of wildlife - threats to biodiversity and human health. Science 287:443.

10. de Castro, F., and B. Bolker. 2005. Mechanisms of disease-induced extinction. Ecology Letters 8:117–126.

11. Direnzo, G. V., P. F. Langhammer, K. R. Zamudio, and K. R. Lips. 2014. Fungal infection intensity and zoospore output of Atelopus zeteki, a potential acute chytrid supershedder. PLoS One 9:e93356.

12. Esteban, J. I., B. Oporto, G. Aduriz, R. A. Juste, and A. Hurtado. 2009. Faecal shedding and strain diversity of Listeria monocytogenes in healthy ruminants and swine in Northern Spain. BMC Vet Res 5:2.

13. Fisher, M. C., D. A. Henk, C. J. Briggs, J. S. Brownstein, L. C. Madoff, S. L. McCraw, and S. J. Gurr. 2012. Emerging fungal threats to animal, plant and ecosystem health. Nature 484:186–194.

14. Frick, W. F., S. J. Puechmaille, J. R. Hoyt, B. A. Nickel, K. E. Langwig, J. T. Foster, K. E. Barlow, T. Bartonicka, D. Feller, A. J. Haarsma, C. Herzog, I. Horacek, J. van der Kooij, B. Mulkens, B. Petrov, R. Reynolds, L. Rodrigues, C. W. Stihler, G. G. Turner, and A. M. Kilpatrick. 2015. Disease alters macroecological patterns of North American bats. Global Ecology and Biogeography 24:741–749.

15. Gervasi, S. S., J. Urbina, J. Hua, T. Chestnut, R. A. Relyea, and A. R. Blaustein. 2013. Experimental evidence for American bullfrog (Lithobates catesbeianus) susceptibility to chytrid fungus (Batrachochytrium dendrobatidis). Ecohealth 10:166–171.

16. Godfrey, S. S. 2013. Networks and the ecology of parasite transmission: A framework for wildlife parasitology. Int J Parasitol Parasites Wildl 2:235–245.

17. Henaux, V., and M. D. Samuel. 2011. Avian influenza shedding patterns in waterfowl: implications for surveillance, environmental transmission, and disease spread. J Wildl Dis 47:566–578.

18. Hicks, A. C., S. Darling, J. Flewelling, R. von Linden, C. U. Meteyer, D. Redell, J. P. White, J. Redell, R. Smith, D. Blehert, N. Rayman, J. R. Hoyt, J. C. Okoniewski, and K. E. Langwig. 2021. Environmental transmission of Pseudogymnoascus destructans to hibernating little brown bats. biorxiv.

19. Hopkins, S. R., J. R. Hoyt, J. P. White, H. M. Kaarakka, J. A. Redell, J. E. DePue, W. H. Scullon, A. M. Kilpatrick, and K. E. Langwig. 2021. Continued preference for suboptimal habitat reduces bat survival with white-nose syndrome. Nat Commun 12:166.

20. Hoyt, J. R., A. M. Kilpatrick, and K. E. Langwig. 2021. Ecology and impacts of white-nose syndrome on bats. Nature Reviews Microbiology:1–15.

21. Hoyt, J. R., K. E. Langwig, J. Okoniewski, W. F. Frick, W. B. Stone, and A. M. Kilpatrick. 2015. Long-term persistence of Pseudogymnoascus destructans, the causative agent of white-nose syndrome, in the absence of bats. EcoHealth 12:330–333.

22. Hoyt, J. R., K. E. Langwig, K. Sun, G. Lu, K. L. Parise, T. Jiang, W. F. Frick, J. T. Foster, J. Feng, and A. M. Kilpatrick. 2016. Host persistence or extinction from emerging infectious disease: insights from white-nose syndrome in endemic and invading regions. Proceedings of the Royal Society B: Biological Sciences 283.

23. Hoyt, J. R., K. E. Langwig, K. Sun, K. L. Parise, A. Li, Y. Wang, X. Huang, L. Worledge, H. Miller, J. P. White, H. M. Kaarakka, J. A. Redell, T. Görföl, S. A. Boldogh, D. Fukui, M. Sakuyama, S. Yachimori, A. Sato, M. Dalannast, A. Jargalsaikhan, N. Batbayar, Y. Yovel, E. Amichai, I. Natradze, W. F. Frick, J. T. Foster, J. Feng, and A. M. Kilpatrick. 2020. Environmental reservoir dynamics predict global infection patterns and population impacts for the fungal disease white-nose syndrome. Proceedings of the National Academy of Sciences:201914794.

24. Hoyt, J. R., K. E. Langwig, J. P. White, H. M. Kaarakka, J. A. Redell, A. Kurta, J. E. DePue, W.H. Scullon, K. L. Parise, and J. T. Foster. 2018. Cryptic connections illuminate pathogen transmission within community networks. Nature:1.

25. Hoyt, J. R., K. L. Parise, J. E. DePue, H. M. Kaarakka, J. A. Redell, W. H. Scullon, R. O’Reskie, J. T. Foster, A. M. Kilpatrick, K. E. Langwig, and J. P. White. 2023. Reducing environmentally mediated transmission to moderate impacts of an emerging wildlife disease. Journal of Applied Ecology 00:1–11.

26. Islam, M. S., M. H. Zaman, M. S. Islam, N. Ahmed, and J. D. Clemens. 2020. Environmental reservoirs of Vibrio cholerae. Vaccine 38 Suppl 1:A52–A62.

27. Jones, K. E., N. G. Patel, M. A. Levy, A. Storeygard, D. Balk, J. L. Gittleman, and P. Daszak. 2008. Global trends in emerging infectious diseases. Nature 451:990–U994.

28. Kailing, M. J., J. R. Hoyt, J. P. White, H. M. Kaarakka, J. A. Redell, A. E. Leon, T. E. Rocke, J. E. DePue, W. H. Scullon, K. L. Parise, J. T. Foster, A. M. Kilpatrick, and K. E. Langwig. 2023. Sex-biased infections scale to population impacts for an emerging wildlife disease. Proc Biol Sci 290:20230040.

29. Kilonzo, C., X. Li, E. J. Vivas, M. T. Jay-Russell, K. L. Fernandez, and E. R. Atwill. 2013. Fecal shedding of zoonotic food-borne pathogens by wild rodents in a major agricultural region of the central California coast. Appl Environ Microbiol 79:6337–6344.

30. Kilpatrick, A. M., P. Daszak, M. J. Jones, P. P. Marra, and L. D. Kramer. 2006. Host heterogeneity dominates West Nile virus transmission. Proceedings of the Royal Society B-Biological Sciences 273:2327–2333.

31. Laggan, N. A., K. L. Parise, J. P. White, H. M. Kaarakka, J. A. Redell, J. E. DePue, W. H. Scullon, J. Kath, J. T. Foster, A. M. Kilpatrick, K. E. Langwig, and J. R. Hoyt. 2023. Data for: Host infection dynamics and disease induced mortality modify species contributions to the environmental reservoir. DRYAD.

32. Langwig, K. E., W. F. Frick, J. T. Bried, A. C. Hicks, T. H. Kunz, and A. M. Kilpatrick. 2012. Sociality, density-dependence and microclimates determine the persistence of populations suffering from a novel fungal disease, white-nose syndrome. Ecol Lett 15.

33. Langwig, K. E., W. F. Frick, J. R. Hoyt, K. L. Parise, K. P. Drees, T. H. Kunz, J. T. Foster, and A. M. Kilpatrick. 2016. Drivers of variation in species impacts for a multi-host fungal disease of bats. Philosophical Transactions of the Royal Society B: Biological Sciences 10.1098/rstb.2015.0456.

34. Langwig, K. E., W. F. Frick, R. Reynolds, K. L. Parise, K. P. Drees, J. R. Hoyt, T. L. Cheng, T. H. Kunz, J. T. Foster, and A. M. Kilpatrick. 2015a. Host and pathogen ecology drive the seasonal dynamics of a fungal disease, white-nose syndrome. Proceedings of the Royal Society B: Biological Sciences 282:20142335.

35. Langwig, K. E., J. R. Hoyt, K. L. Parise, W. F. Frick, J. T. Foster, and A. M. Kilpatrick. 2017. Resistance in persisting bat populations after white-nose syndrome invasion. Philosophical Transactions of the Royal Society B: Biological Sciences 372:20160044.

36. Langwig, K. E., J. R. Hoyt, K. L. Parise, J. Kath, D. Kirk, W. F. Frick, J. T. Foster, and A. M. Kilpatrick. 2015b. Invasion Dynamics of White-Nose Syndrome Fungus, Midwestern United States, 2012-2014. Emerging Infectious Disease 21:1023–1026.

37. Langwig, K. E., J. P. White, K. L. Parise, H. M. Kaarakka, J. A. Redell, J. E. DePue, W. H. Scullon, J. T. Foster, A. M. Kilpatrick, and J. R. Hoyt. 2021. Mobility and infectiousness in the spatial spread of an emerging fungal pathogen. J Anim Ecol 90:1134–1141.

38. Lawley, T. D., D. M. Bouley, Y. E. Hoy, C. Gerke, D. A. Relman, and D. M. Monack. 2008. Host transmission of Salmonella enterica serovar Typhimurium is controlled by virulence factors and indigenous intestinal microbiota. Infect Immun 76:403–416.

39. Lenth, R. V. 2022. emmeans: Estimated Marginal Means, aka Least-Squares Means.

40. Lloyd-Smith, J. O., S. J. Schreiber, P. E. Kopp, and W. M. Getz. 2005. Superspreading and the effect of individual variation on disease emergence. Nature 438:355–359.

41. Lockwood, J. L., P. Cassey, and T. Blackburn. 2005. The role of propagule pressure in explaining species invasions. Trends Ecol Evol 20:223–228.

42. Lorch, J. M., C. U. Meteyer, M. J. Behr, J. G. Boyles, P. M. Cryan, A. C. Hicks, A. E. Ballmann, J. T. H. Coleman, D. N. Redell, D. M. Reeder, and D. S. Blehert. 2011. Experimental infection of bats with Geomyces destructans causes white-nose syndrome. Nature 480:376–378.

43. Lorch, J. M., L. K. Muller, R. E. Russell, M. O’Connor, D. L. Lindner, and D. S. Blehert. 2013. Distribution and environmental persistence of the causative agent of white-nose syndrome, Geomyces destructans, in bat hibernacula of the eastern United States. Applied and Environmental Microbiology 79:1293–1301.

44. Maguire, C., G. V. DiRenzo, T. S. Tunstall, C. R. Muletz-Wolz, K. R. Zamudio, and K. R. Lips. 2016. Dead or alive? Viability of chytrid zoospores shed from live amphibian hosts. Diseases of Aquatic Organisms 199:179–187.

45. Matthews, L., J. C. Low, D. L. Gally, M. C. Pearce, D. J. Mellor, J. A. Heesterbeek, M. Chase-Topping, S. W. Naylor, D. J. Shaw, S. W. Reid, G. J. Gunn, and M. E. Woolhouse. 2006. Heterogeneous shedding of Escherichia coli O157 in cattle and its implications for control. Proc Natl Acad Sci U S A 103:547–552.

46. McGuire, L. P., H. W. Mayberry, and C. K. Willis. 2017. White-nose syndrome increases torpid metabolic rate and evaporative water loss in hibernating bats. American Journal of Physiology-Regulatory, Integrative and Comparative Physiology 313:R680–R686.

47. Mitchell, K. M., T. S. Churcher, T. W. Garner, and M. C. Fisher. 2008. Persistence of the emerging pathogen Batrachochytrium dendrobatidis outside the amphibian host greatly increases the probability of host extinction. Proc Biol Sci 275:329–334.

48. Muller, L. K., J. M. Lorch, D. L. Lindner, M. O’Connor, A. Gargas, and D. S. Blehert. 2013. Bat white-nose syndrome: a real-time TaqMan polymerase chain reaction test targeting the intergenic spacer region of Geomyces destructans. Mycologia 105:253–259.

49. Munywoki, P. K., D. C. Koech, C. N. Agoti, N. Kibirige, J. Kipkoech, P. A. Cane, G. F. Medley, and D. J. Nokes. 2015. Influence of age, severity of infection, and co-infection on the duration of respiratory syncytial virus (RSV) shedding. Epidemiol Infect 143:804–812.

50. Paull, S. H., S. Song, K. M. McClure, L. C. Sackett, A. M. Kilpatrick, and P. T. J. Johnson. 2012. From superspreaders to disease hotspots: linking transmission across hosts and space. Frontiers in Ecology and the Environment 10:75–82.

51. Peterson, A. C., and V. J. McKenzie. 2014. Investigating differences across host species and scales to explain the distribution of the amphibian pathogen Batrachochytrium dendrobatidis. PLoS One 9:e107441.

52. Plummer, I. H., C. J. Johnson, A. R. Chesney, J. A. Pedersen, and M. D. Samuel. 2018. Mineral licks as environmental reservoirs of chronic wasting disease prions. PLoS One 13:e0196745.

53. Rushmore, J., D. Caillaud, L. Matamba, R. M. Stumpf, S. P. Borgatti, and S. Altizer. 2013. Social network analysis of wild chimpanzees provides insights for predicting infectious disease risk. J Anim Ecol 82:976–986.

54. Scheele, B. C., D. A. Hunter, L. A. Brannelly, L. F. Skerratt, and D. A. Driscoll. 2017. Reservoir-host amplification of disease impact in an endangered amphibian. Conserv Biol 31:592–600.

55. Searle, C. L., S. S. Gervasi, J. Hua, J. I. Hammond, R. A. Relyea, D. H. Olson, and A. R. Blaustein. 2011. Differential Host Susceptibility to Batrachochytrium dendrobatidis, an Emerging Amphibian Pathogen. Conservation Biology 25:965–974.

56. Searle, S. R., F. M. Speed, and G. A. Miliken. 1980. Population Marginal Means in the Linear Model: An Alternative to Least Squares Means. The American Statistician 34:216–221.

57. Sheth, P. M., A. Danesh, A. Sheung, A. Rebbapragada, K. Shahabi, C. Kovacs, R. Halpenny, D. Tilley, T. Mazzulli, K. MacDonald, D. Kelvin, and R. Kaul. 2006. Disproportionately high semen shedding of HIV is associated with compartmentalized cytomegalovirus reactivation. J Infect Dis 193:45–48.

58. Slater, N., R. M. Mitchell, R. H. Whitlock, T. Fyock, A. K. Pradhan, E. Knupfer, Y. H. Schukken, and Y. Louzoun. 2016. Impact of the shedding level on transmission of persistent infections in Mycobacterium avium subspecies paratuberculosis (MAP). Vet Res 47:38.

59. Taylor, L. H., S. M. Latham, and M. E. Woolhouse. 2001. Risk factors for human disease emergence. Philos Trans R Soc Lond B Biol Sci 356:983–989.

60. Turner, W. C., K. L. Kausrud, W. Beyer, W. R. Easterday, Z. R. Barandongo, E. Blaschke, C. C. Cloete, J. Lazak, M. N. Van Ert, and H. H. Ganz. 2016. Lethal exposure: An integrated approach to pathogen transmission via environmental reservoirs. Scientific reports 6:27311.

61. VanderWaal, K. L., and V. O. Ezenwa. 2016. Heterogeneity in pathogen transmission: mechanisms and methodology. Funct. Ecol. 30:1606–1622.

62. Verant, M. L., M. U. Carol, J. R. Speakman, P. M. Cryan, J. M. Lorch, and D. S. Blehert. 2014. White-nose syndrome initiates a cascade of physiologic disturbances in the hibernating bat host. BMC physiology 14:10.

63. Warnecke, L., J. M. Turner, T. K. Bollinger, J. M. Lorch, V. Misra, P. M. Cryan, G. Wibbelt, D. S. Blehert, and C. K. R. Willis. 2012. Inoculation of bats with European Geomyces destructans supports the novel pathogen hypothesis for the origin of white-nose syndrome. Proceedings of the National Academy of Sciences of the United States of America 109:6999–7003.

64. Warnecke, L., J. M. Turner, T. K. Bollinger, V. Misra, P. M. Cryan, D. S. Blehert, G. Wibbelt, and C. K. R. Willis. 2013. Pathophysiology of white-nose syndrome in bats: a mechanistic model linking wing damage to mortality. Biology Letters 9.

