## Appendix S1 for "Host infection dynamics and disease induced mortality modify species contributions to the environmental reservoir"

**Journal name:** Ecology
**Title:** Host infection dynamics and disease induced mortality modify species contributions to the environmental reservoir

| **Citation in manuscript** | **Description** |
|---|---|
| Appendix S1 | First appendix |
| Figure S1 | Yearly species abundance across sites |
| Figure S2 | Correlation between qPCR and CFU's |
| Figure S3 | Coefficient of variation for within species pathogen shedding and infection intensity during invading and established stages |
| Figure S4 | Relationship between species abundance and site-level contamination |
| Figure S5 | Coefficients of the slopes for pathogen shedding as a function of infection intensity |
| Table S1 | Table of years of sampling for each site by invasion stage |
| Table S2 | Table with sample sizes |
| Table S3 | Table describing statistical approach |
| Table S4 | Statistical table for Fig1A |
| Table S5 | Statistical table for Fig1B |
| Table S6 | Statistical table for Fig 2A |
| Table S7 | Statistical table for Fig 2B |
| Table S8 | Statistical table for Fig 2C |
| Table S9 | Statistical table for Fig 3 (invading) |
| Table S10 | Statistical table for Fig 3 (established) |
| Table S11 | Statistical table for Fig 4A |
| Table S12 | Statistical table for Fig 4B |
| Table S13 | Statistical table for Fig S4 |

### Supplementary Figures

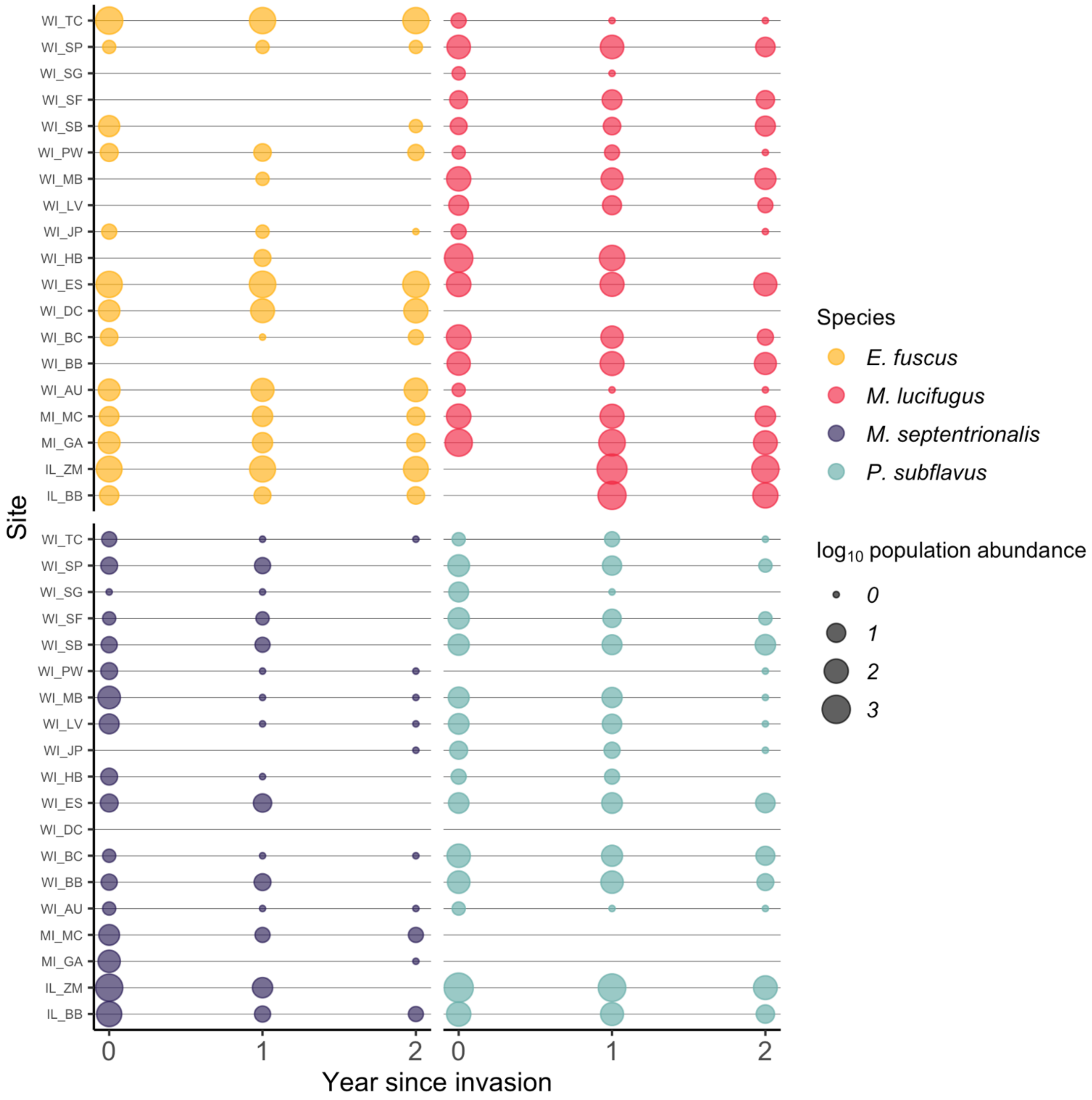

**Figure S1. Species $\log_{10}$ population abundance for each site by year (years 0-2)**. Points indicate the $\log_{10}$ population abundance of each species within a site for each year of data collection. Different species are displayed by color and point size indicates $\log_{10}$ population abundance within each site.

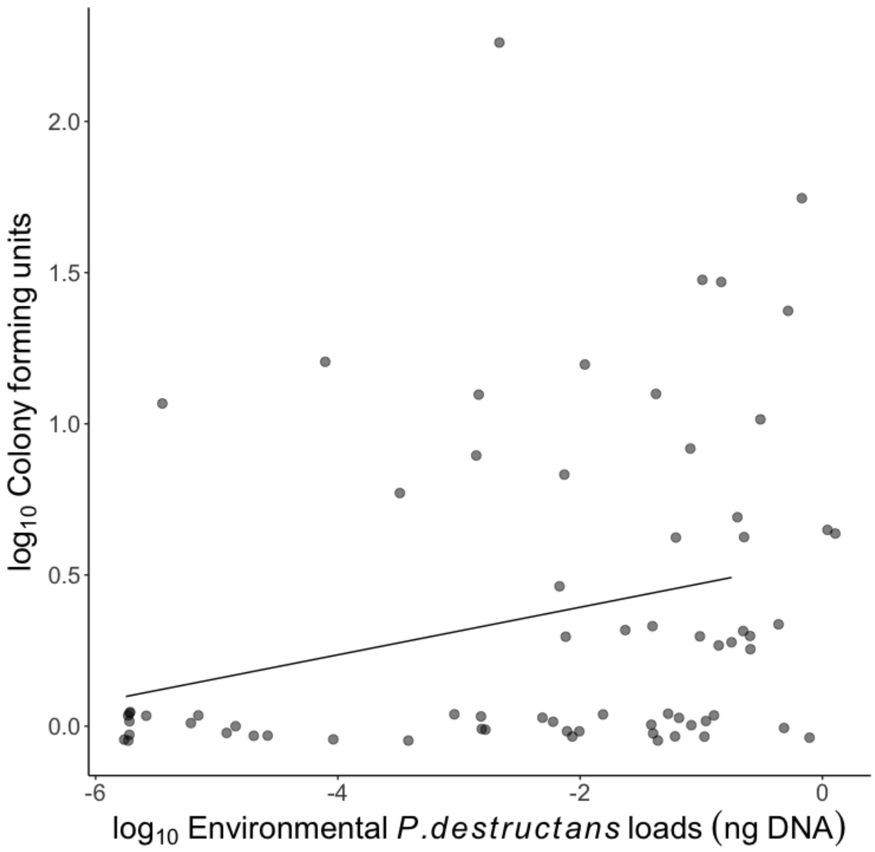

**Figure S2. Relationship between viable colony forming units and fungal loads measured using qPCR in environmental swabs.** The number of viable fungal colonies increased significantly with fungal loads using qPCR (CFU: intercept = 0.55 ± 0.1, slope = 0.08, relationship between CFU's and environmental fungal loads $P < 0.02$).

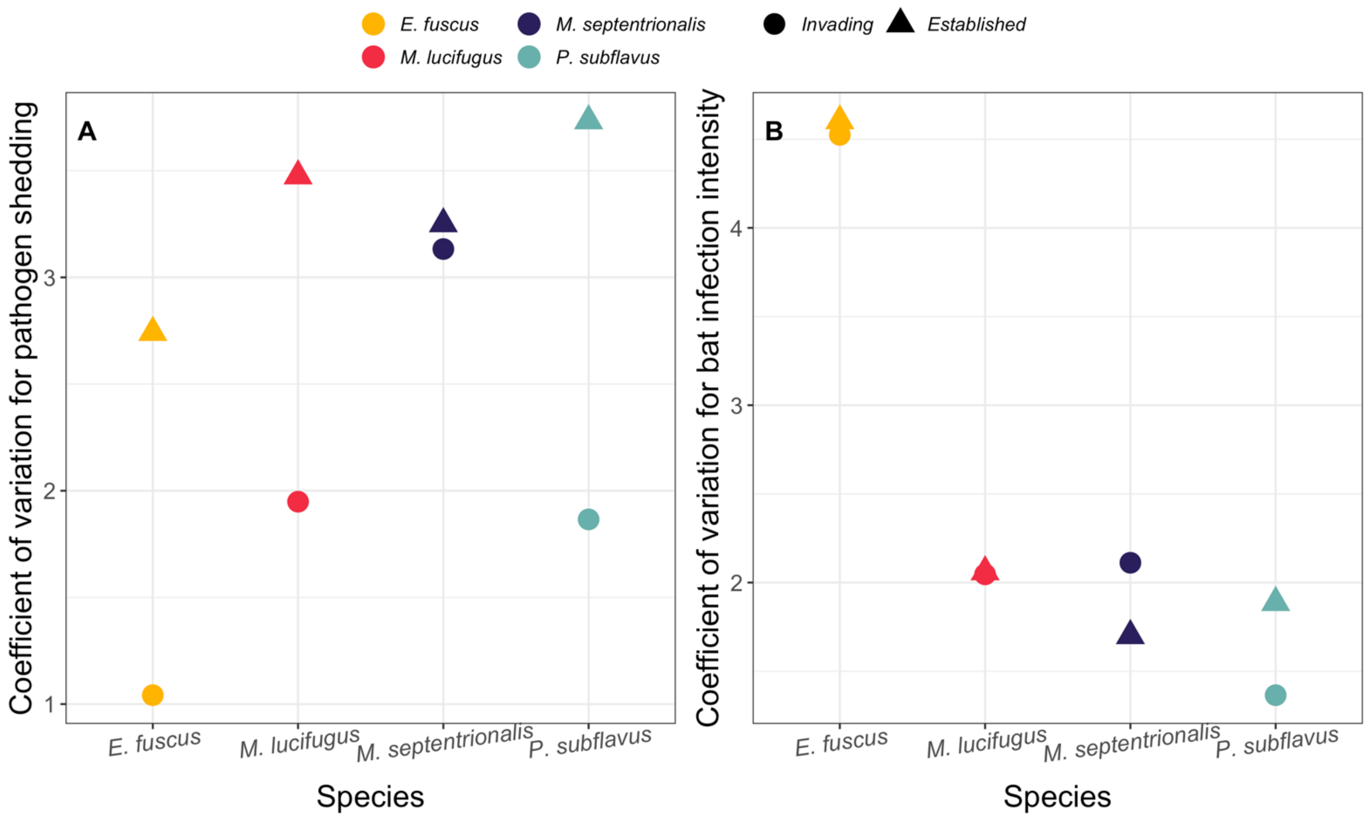

**Figure S3. Coefficient of variation for within species pathogen shedding and infection intensity during invading and established stages.** Each point represents the coefficient of variation within species populations. (A) Coefficient of variation of *P. destructans* environmental loads (ng DNA) within species during pathogen invasion and establishment in late hibernation. (B) Coefficient of variation of *P. destructans* bat loads (ng DNA) within species during pathogen invasion and establishment in late hibernation.

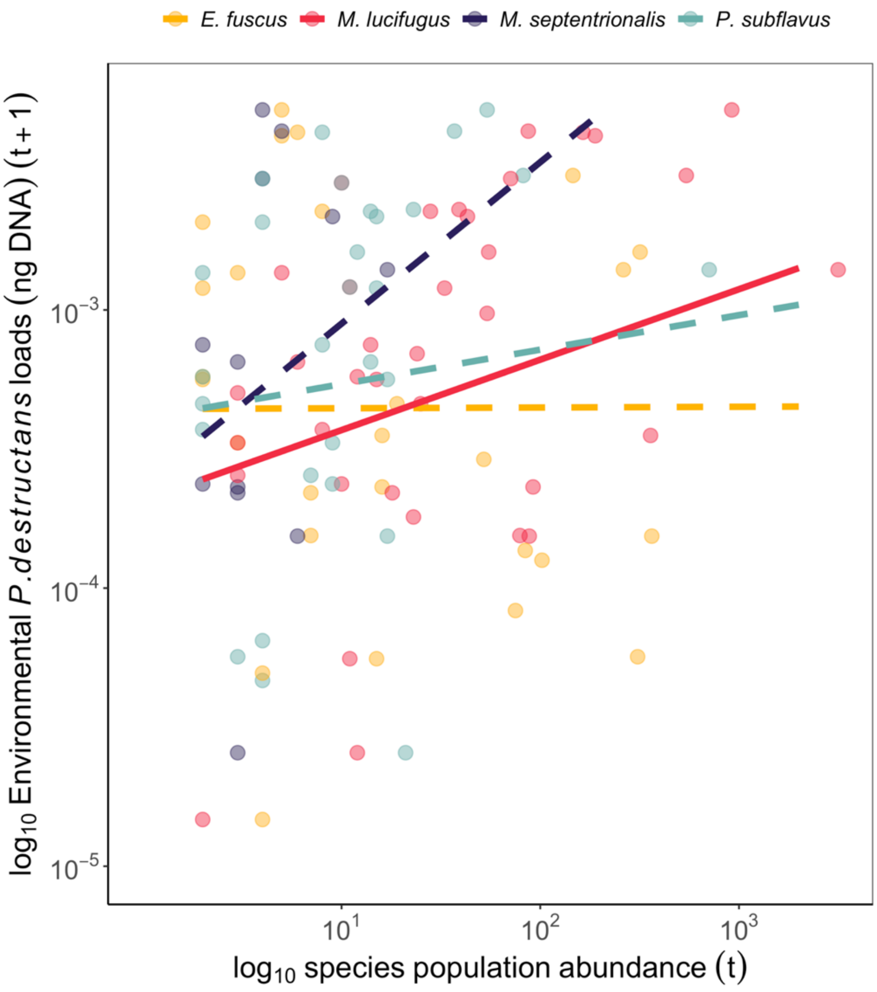

**Figure S4. Effects of host abundance on site level environmental contamination.** Species within the community are differentiated by color and points represent a species population at an individual site. Total $\log_{10}$ species abundance in early hibernation predicting $\log_{10}$ environmental *P. destructans* loads (ng DNA) taken from samples > 2m away from roosting bats within sites during late hibernation during the established stage. Solid lines indicate support for a positive relationship between species abundance and site-level environmental contamination (Table S7). Points for *E. fuscus* population at abundance $10^{2.47}$, and environmental loads $10^{-5.22}$ are not shown on figure for scaling purposes.

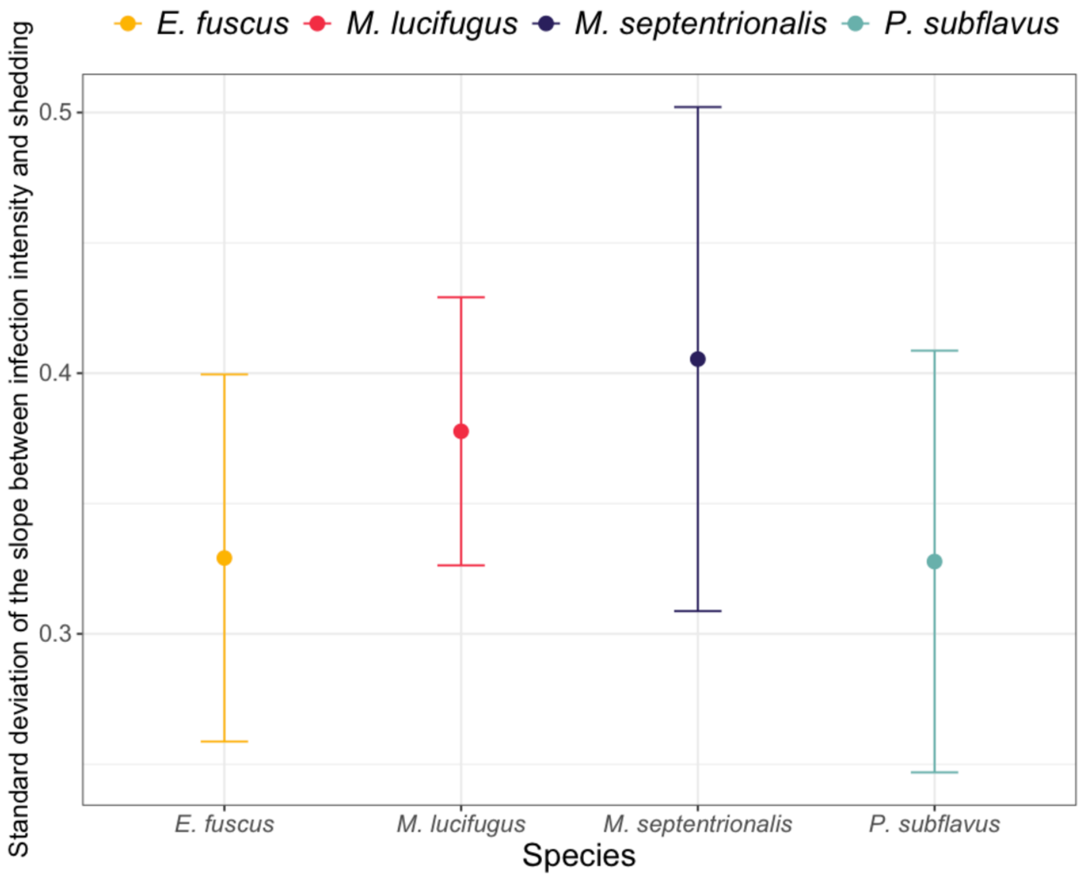

**Figure S5. Coefficients of the slopes for pathogen shedding as a function of infection intensity do not differ among species.** Slope coefficients and standard error for the relationship between infection intensity and pathogen shedding by species. The standard error of the relationship between $\log_{10}$ *P. destructans* loads (ng DNA) on an individual bat and the amount of $\log_{10}$ *P. destructans* (ng DNA) in the environment directly underneath each individual during pathogen invasion and establishment in late hibernation. Points represent the standard deviation and bars indicate ± standard error for each species.

**Supplemental tables**

**Table S1. Years of sampling for each site by invasion stage**. Invading years are the first year the pathogen arrived and established years are the following two years after invasion.

| Site | Invading year | Established years |
|---|---|---|
| WI_AU | 2017 | 2018-2019 |
| WI_BC | 2016 | 2017-2018 |
| IL_BB | 2013 | 2014-2015 |
| WI_BB | 2015 | 2016-2017 |
| WI_DC | 2017 | 2018-2019 |
| WI_ES | 2016 | 2017-2018 |
| MI_GA | 2015 | 2016-2017 |
| WI_HB | 2015 | 2016-2017 |
| WI_JP | 2015 | 2016-2017 |
| WI_LV | 2017 | 2018-2019 |
| WI_MB | 2016 | 2017-2018 |
| MI_MC | 2015 | 2016-2017 |
| WI_PW | 2016 | 2017-2018 |
| WI_SP | 2017 | 2018-2019 |
| WI_SF | 2015 | 2016-2017 |
| WI_SG | 2015 | 2016-2017 |
| WI_SB | 2015 | 2016-2017 |
| WI_TC | 2017 | 2018-2019 |
| IL_ZM | 2013 | 2014-2015 |

**Table S2. Sample sizes by site for each species by invasion stage**. Environmental samples were samples taken from substrate directly underneath a roosting bat on the wall or ceiling during either invading or established stages. Bat infection samples were epidermal swab samples taken from hibernating bats.

| Site | Species | Environmental samples during invading | Environmental samples during established | Bat infection samples |
|---|---|---|---|---|
| WI_AU | *E. fuscus* | 6 | 17 | 23 |
| WI_AU | *M. lucifugus* | 1 | 0 | 1 |
| WI_BC | *E. fuscus* | 2 | 1 | 3 |
| WI_BC | *M. lucifugus* | 7 | 9 | 16 |
| WI_BC | *M. septentrionalis* | 1 | 0 | 1 |
| WI_BC | *P. subflavus* | 7 | 15 | 22 |
| IL_BB | *E. fuscus* | 4 | 4 | 8 |
| IL_BB | *M. lucifugus* | 6 | 20 | 26 |
| IL_BB | *M. septentrionalis* | 4 | 3 | 7 |
| IL_BB | *P. subflavus* | 5 | 19 | 24 |
| WI_BB | *M. lucifugus* | 7 | 15 | 22 |
| WI_BB | *M. septentrionalis* | 2 | 2 | 4 |
| WI_BB | *P. subflavus* | 9 | 11 | 20 |
| WI_DC | *E. fuscus* | 7 | 19 | 26 |
| WI_ES | *E. fuscus* | 7 | 18 | 25 |
| WI_ES | *M. lucifugus* | 7 | 19 | 26 |
| WI_ES | *M. septentrionalis* | 5 | 7 | 12 |
| WI_ES | *P. subflavus* | 7 | 18 | 25 |
| MI_GA | *E. fuscus* | 0 | 10 | 10 |
| MI_GA | *M. lucifugus* | 3 | 15 | 18 |
| MI_GA | *M. septentrionalis* | 7 | 0 | 7 |
| WI_HB | *E. fuscus* | 0 | 2 | 2 |
| WI_HB | *M. lucifugus* | 11 | 10 | 21 |
| WI_HB | *M. septentrionalis* | 4 | 0 | 4 |
| WI_HB | *P. subflavus* | 1 | 5 | 6 |
| WI_JP | *E. fuscus* | 2 | 1 | 3 |
| WI_JP | *M. lucifugus* | 1 | 0 | 1 |
| WI_JP | *P. subflavus* | 2 | 3 | 5 |
| WI_LV | *M. lucifugus* | 6 | 9 | 15 |
| WI_LV | *M. septentrionalis* | 6 | 0 | 6 |
| WI_LV | *P. subflavus* | 7 | 7 | 14 |
| WI_MB | *E. fuscus* | 0 | 1 | 1 |
| WI_MB | *M. lucifugus* | 7 | 17 | 24 |
| WI_MB | *M. septentrionalis* | 7 | 0 | 7 |

| | | | | |
|---|---|---|---|---|
| WI_MB | *P. subflavus* | 7 | 6 | 13 |
| MI_MC | *E. fuscus* | 5 | 12 | 17 |
| MI_MC | *M. lucifugus* | 4 | 16 | 20 |
| MI_MC | *M. septentrionalis* | 6 | 5 | 11 |
| WI_PW | *E. fuscus* | 3 | 6 | 9 |
| WI_PW | *M. lucifugus* | 1 | 0 | 1 |
| WI_PW | *M. septentrionalis* | 3 | 0 | 3 |
| WI_SP | *E. fuscus* | 1 | 1 | 2 |
| WI_SP | *M. lucifugus* | 7 | 20 | 27 |
| WI_SP | *M. septentrionalis* | 2 | 2 | 4 |
| WI_SP | *P. subflavus* | 9 | 8 | 17 |
| WI_SF | *M. lucifugus* | 3 | 9 | 12 |
| WI_SF | *M. septentrionalis* | 1 | 1 | 2 |
| WI_SF | *P. subflavus* | 9 | 5 | 14 |
| WI_SG | *M. lucifugus* | 1 | 0 | 1 |
| WI_SG | *P. subflavus* | 7 | 0 | 7 |
| WI_SB | *E. fuscus* | 8 | 1 | 9 |
| WI_SB | *M. lucifugus* | 2 | 14 | 16 |
| WI_SB | *M. septentrionalis* | 1 | 1 | 2 |
| WI_SB | *P. subflavus* | 9 | 15 | 24 |
| WI_TC | *E. fuscus* | 8 | 19 | 27 |
| WI_TC | *P. subflavus* | 0 | 1 | 1 |
| IL_ZM | *E. fuscus* | 5 | 20 | 25 |
| IL_ZM | *M. lucifugus* | 5 | 21 | 26 |
| IL_ZM | *M. septentrionalis* | 3 | 1 | 4 |
| IL_ZM | *P. subflavus* | 4 | 20 | 24 |

**Table S3: Description of statistical approach and reporting for analyses.**

| Figure | Analysis | Reference level | Reporting |
|---|---|---|---|
| 1A | Pathogen shedding by species during the invading year | *M. septentrionalis* | lmer model output |
| 1B | Pathogen shedding by species during established stage | *M. lucifugus* | lmer model output |
| 2A | Infection prevalence by species | *M. lucifugus* | glmer model output |
| 2B | Infection intensity by species | *E. fuscus* | lmer model output |
| 2C | The relationship between infection intensity and pathogen shedding by species | N/A | Estimated marginal means of linear trends |
| 3 | Species population abundance during the invading year | *M. lucifugus* | lmer model output |
| 3 | Species population abundance during the established stage | *M. lucifugus* | lmer model output |
| 4A | Species propagule pressure during the invading year | *M. septentrionalis* | glmer model output |
| 4B | Species propagule pressure during the established stage | N/A | Estimated marginal means |
| 4C | Propagule pressure predict site-level contamination | N/A | lmer model output |
| S2 | Species population abundance predicting site-level pathogen contamination during the established stage | *M. lucifugus* | lmer model output |

**Table S4. Coefficients from linear mixed effects model for pathogen shedding by species during the invading stage.**

Environmental fungal loads was the response variable, species was a fixed effect and site was a random effect during late hibernation. Results are visualized in Fig. 1A.

| | Estimate | Standard error | Degrees of freedom | t value | Pr(>\|t\|) |
|---|---|---|---|---|---|
| Intercept (*M. septentrionalis*) | -3.72 | 0.23 | 33.91 | -16.22 | 0.00 |
| *E. fuscus* | -0.88 | 0.33 | 73.60 | -2.68 | 0.01 |
| *M. lucifugus* | -0.52 | 0.22 | 69.50 | -2.30 | 0.02 |
| *P. subflavus* | -0.86 | 0.35 | 73.99 | -2.48 | 0.02 |

**Table S5. Coefficients from linear mixed effects model for pathogen shedding by species during the established stage.**

Environmental fungal loads was the response variable, species was a fixed effect and site was a random effect during late hibernation. Results are visualized in Fig. 1B.

| | Estimate | Standard error | Degrees of freedom | t value | Pr(>\|t\|) |
|---|---|---|---|---|---|
| Intercept (*M. lucifugus*) | -3.41 | 0.12 | 27.54 | -27.54 | 0.00 |
| *E. fuscus* | -0.75 | 0.14 | 297.01 | -5.29 | 0.00 |
| *M. septentrionalis* | -0.33 | 0.24 | 362.70 | -1.39 | 0.17 |
| *P. subflavus* | -0.65 | 0.13 | 371.04 | -5.01 | 0.00 |

**Table S6. Coefficients from generalized linear mixed effects model for infection prevalence by species.**

We fit a generalized linear mixed effects model with a binomial distribution, with bat infection prevalence as the response variable, species as a fixed effect, and site as a random effect during late hibernation for both invasion and established stage (years 0-2). Results are visualized in Fig. 2A.

| | Estimate | Standard Error | z value | Pr(>\|z\|) |
|---|---|---|---|---|
| Intercept (*M. lucifugus*) | 2.19 | 0.27 | 8.06 | 0.00 |
| *E. fuscus* | -0.79 | 0.31 | -2.57 | 0.01 |
| *M. septentrionalis* | -0.83 | 0.35 | -2.38 | 0.02 |
| *P. subflavus* | -1.10 | 0.24 | -4.50 | 0.00 |

**Table S7. Coefficients from linear mixed effects model for infection intensity by species.**

Bat fungal loads was the response variable, species was a fixed effect, and site was a random effect during late hibernation for invasion and established stage (years 0-2). Results are visualized in Fig. 2B.

| | Estimate | Standard Error | Degrees of Freedom | t value | Pr(>\|t\|) |
|---|---|---|---|---|---|
| Intercept (*E. fuscus*) | -3.37 | 0.16 | 50.90 | -21.49 | 0.00 |
| *M. lucifugus* | 1.38 | 0.15 | 440.31 | 8.90 | 0.00 |
| *M. septentrionalis* | 1.08 | 0.21 | 581.98 | 5.10 | 0.00 |
| *P. subflavus* | 1.30 | 0.17 | 418.62 | 7.68 | 0.00 |

**Table S8. Estimated marginal means of linear trends from a linear mixed effects model of pathogen shedding predicted by species infections.**

Environmental fungal loads was the response variable, species interacting with and bat infection intensity were fixed effect predictors, and site was a random effect, during late hibernation for invasion and established years (years 0-2). Results are visualized in Fig. 2C.

| Estimated marginal means of linear trends | | | | | |
|---|---|---|---|---|---|
| | Slope trend | Standard error | Degrees of freedom | t ratio | p value |
| *E. fuscus* | 0.33 | 0.07 | 597 | 4.66 | 0.00 |
| *M. lucifugus* | 0.38 | 0.05 | 593 | 7.31 | 0.00 |
| *M. septentrionalis* | 0.41 | 0.10 | 595 | 4.18 | 0.00 |
| *P. subflavus* | 0.33 | 0.08 | 594 | 4.05 | 0.00 |
| Contrast | Estimate | Standard error | Degrees of freedom | t ratio | p value |
| (*E. fuscus* slope) – (*M. lucifugus* slope) | -0.05 | 0.09 | 596 | -0.56 | 0.94 |
| (*E. fuscus* slope) – (*M. septentrionalis* slope) | -0.08 | 0.12 | 597 | -0.63 | 0.92 |
| (*E. fuscus* slope) – (*P. subflavus* slope) | 0.00 | 0.11 | 592 | 0.01 | 1.00 |
| (*M. lucifugus* slope) – (*M. septentrionalis* slope) | -0.03 | 0.11 | 585 | -0.26 | 0.99 |
| (*M. lucifugus* slope) – (*P. subflavus* slope) | 0.05 | 0.10 | 596 | 0.52 | 0.95 |
| (*M. septentrionalis* slope) – (*P. subflavus* slope) | 0.08 | 0.13 | 595 | 0.61 | 0.93 |

**Table S9. Coefficients from linear mixed effects model for population abundance by species during invading stage.**

$\log_{10}$ Population abundance was the response variable, species was a fixed effect, and site was a random effect. Results are visualized in Fig. 3.

| | Estimate | Standard Error | Degrees of Freedom | t value | Pr(>\|t\|) |
|---|---|---|---|---|---|
| Intercept (*M. lucifugus)* | 1.63 | 0.19 | 53.02 | 8.44 | 0.00 |
| *E. fuscus* | -0.65 | 0.24 | 43.07 | -2.74 | 0.01 |
| *M. septentrionalis* | -0.60 | 0.21 | 40.71 | -2.81 | 0.01 |
| *P. subflavus* | -0.41 | 0.23 | 42.89 | -1.80 | 0.08 |

**Table S10. Coefficients from linear mixed effects model for population abundance by species during established stage.**

$\log_{10}$ Population abundance was the response variable, species was a fixed effect, and site was a random effect. Results are visualized in Fig. 3.

| | Estimate | Standard Error | Degrees of Freedom | t value | Pr(>\|t\|) |
|---|---|---|---|---|---|
| Intercept (*M. lucifugus*) | 1.28 | 0.13 | 66.04 | 9.77 | 0.00 |
| *E. fuscus* | -0.54 | 0.16 | 122.91 | -3.34 | 0.00 |
| *M. septentrionalis* | -0.76 | 0.14 | 114.72 | -5.33 | 0.00 |
| *P. subflavus* | -0.38 | 0.15 | 118.83 | -2.58 | 0.01 |

**Table S11. Coefficients from generalized linear mixed effects model with negative binomial distribution for propagule pressure by species during invading stage.**

Propagule pressure was the response variable, species was a fixed effect, and site was a random effect during invasion (year 0). Results are visualized in Fig. 4A.

| | Estimate | Standard Error | z value | Pr(>\|z\|) |
|---|---|---|---|---|
| Intercept (*M. septentrionalis*) | 7.44 | 1.05 | 7.09 | 0.00 |
| *E. fuscus* | -4.27 | 1.21 | -3.54 | 0.00 |
| *M. lucifugus* | 0.87 | 1.03 | 0.85 | 0.39 |
| *P. subflavus* | -3.17 | 1.25 | -2.55 | 0.01 |

**Table S12. Estimated marginal means from generalized linear mixed effects model with negative binomial distribution for propagule pressure by species during established stage.**

Propagule pressure was the response variable, species was a fixed effect, and site was a random effect during establishment (years 1-2). Results are visualized in Fig. 4B.

| Species | Estimated marginal mean | Standard error | Degrees of freedom | Lower confidence limit | Upper confidence limit |
|---|---|---|---|---|---|
| *E. fuscus* | 6.91 | 0.54 | 93 | 5.83 | 7.98 |
| *M. lucifugus* | 10.75 | 0.51 | 93 | 9.74 | 11.76 |
| *M. septentrionalis* | 6.52 | 0.81 | 93 | 4.90 | 8.13 |
| *P. subflavus* | 8.3 | 0.54 | 93 | 7.23 | 9.36 |
| Contrast | Estimate | Standard error | Degrees of freedom | t ratio | p value |
| *E. fuscus* – *M. lucifugus* | -3.84 | 0.56 | 93 | -6.86 | 0.00 |
| *E. fuscus* – *M. septentrionalis* | 0.39 | 0.83 | 93 | 0.47 | 0.97 |
| *E. fuscus* – *P. subflavus* | -1.39 | 0.63 | 93 | -2.21 | 0.13 |
| *M. lucifugus* – *M. septentrionalis* | 4.24 | 0.79 | 93 | 5.34 | 0.00 |
| *M. lucifugus* – *P. subflavus* | 2.46 | 0.54 | 93 | 4.52 | 0.00 |
| *M. septentrionalis* – *P. subflavus* | -1.78 | 0.85 | 93 | -2.09 | 0.16 |

**Table S13. Coefficients from linear mixed effects model for site-level environmental contamination by species.**

Site-level average environmental fungal load was the response variable, $\log_{10}$ population abundance interacting with species were predictor variables, and site was a random effect. Results are visualized in Fig. S4.

| | Estimate | Standard Error | Degrees of Freedom | t value | Pr(>\|t\|) |
|---|---|---|---|---|---|
| Intercept (*M. lucifugus*) | -3.69 | 0.22 | 86.30 | -17.04 | 0.00 |
| log10 abundance | 0.25 | 0.11 | 83.39 | 2.25 | 0.03 |
| *E. fuscus* | 0.33 | 0.23 | 77.79 | 1.41 | 0.16 |
| *M. septentrionalis* | 0.06 | 0.35 | 74.45 | 0.16 | 0.88 |
| *P. subflavus* | 0.29 | 0.24 | 76.76 | 1.24 | 0.22 |
| log10 abundance:*E. fuscus* | -0.25 | 0.15 | 80.67 | -1.63 | 0.11 |
| log10 abundance:*M. septentrionalis* | 0.33 | 0.48 | 74.92 | 0.68 | 0.50 |
| log10 abundance:*P. subflavus* | -0.13 | 0.18 | 76.06 | -0.73 | 0.47 |
